# Visitor effect on the behavior of chimpanzees (*pan troglodyte*) at a primate rescue center

**DOI:** 10.1101/2024.09.21.614280

**Authors:** Sofia Alegre Maurer, Miriam Ross, Jose Gil Dolz

## Abstract

1.

The objective of this study was to examine the impact of visitors on the behavior of chimpanzees at a sanctuary. We hypothesized there wouldn’t be an increase in abnormal, agnostic, or self-directed behaviors during visits, nor a decrease in affiliative behaviors when visitors are present.

The study examined the effects of visitor presence on chimpanzee behavior at Fundació MONA, a rescue center.

**Key findings include:**

- No significant changes in abnormal or self-directed behaviors or agonistic behaviors were observed with visitor presence.
- Affiliative behaviors (excluding grooming) showed a slight decrease during visits.

These results indicate that guided visitor interactions do not adversely affect chimpanzee behavior and may even enhance their welfare. The study supports the implementation of structured visitor programs for public education and funding, without compromising animal well-being. This contrasts with some previous research in zoo settings, suggesting that controlled visits can be beneficial in primate sanctuaries.

## 2. Introduction

Chimpanzees (Pan troglodytes) are a species of social primates native to central and western Africa^1^. In their natural habitat they form socially complex and hierarchical groups of up to 150 individuals led by a dominant male^2,3^. Chimpanzees are currently in danger of extinction^4^. The main threats to the conservation of chimpanzees include illegal hunting for their meat^5^, their capture for the pet trade^6^, their exploitation in entertainment shows^7^, as well as the loss or degradation of their habitat^8^. As a result, there are centers dedicated to the rescue and rehabilitation of chimpanzees that have been mistreated or used as pets or for entertainment shows^9^. These centers offer chimpanzees adequate conditions so that they can develop their natural behaviors and live with others of their species in a social environment^10^.

Most of these rescue centers are private initiatives that are maintained thanks to specific donations or through the visits they offer to learn about the history of these animals and raise awareness among the population about the psychological and physical consequences that these chimpanzees experience after having suffered in the past because of those practices^11^. In recent years, more efforts have been invested to improve the well-being of animals in captivity, both in rescue centers and zoos^12,13^. This concern includes the impact that visitors have on the behavior and well-being of the animals^14,15^.

This was the objective of this study, to evaluate the impact of visitors on the behavior of chimpanzees at a rescue center.

Other investigations across various species of primates have found differing levels of stress with visitors, with chimpanzees appearing to be more prone to stress behaviors than bonobos or macaques.

A similar study to ours looked at bonobos in a zoo and found that the number of visitors didn’t affect their behavior^16^. The bonobos didn’t show more stress or less social interaction when more visitors were around. It’s possible that the bonobos at Twycross Zoo are used to visitors, so their well-being isn’t impacted. Like at MONA, the enclosure design, which lets the bonobos hide from visitors, may help them tolerate having visitors around. However, the researchers noted that visitor impact can vary between different species and even among individuals of the same species.

Most animals get stressed by constant interaction with visitors, but captive bonobos might actually enjoy it, suggesting it might be positive^17^. This study observed bonobos at The Cincinnati Zoo to see if they showed more abnormal behaviors indoors, where they had more visitor interaction, compared to outdoors with less interaction. After 54 hours of observation, there was no significant difference in abnormal behaviors between the two settings, and no link to the number of visitors. This suggests that visitor interactions don’t stress out bonobos, and in fact, might even be good for them. It would be interesting to explore further with more data to see if bonobos really do handle stress better than chimpanzees due to their friendlier nature.

Some studies show primates react differently to visitors—some find them stressful, others enjoy it, and some aren’t affected^18^. Researchers tested four main ideas to explain this: species differences, visitor behavior, enclosure type, and individual traits. They found that small, tree-dwelling primates react more negatively, especially with loud noise, while those in walk-through enclosures usually have positive or neutral reactions. Age and sex didn’t have a clear impact. A mix of factors seems to influence how primates respond, and better tools, like using AI analysis, could help researchers understand these responses better.

A study focusing on lion-tailed macaques shows how visitor presence and behavior can alter the activity and stress-related behaviors in captive primates^19^. Visitors increased stress-related behaviors and altered the daily activity patterns of the macaques.

A study was focused on gorillas, but it also examined how bonobos and other primates responded to visitor crowd sizes^20^. At Apenheul Primate Park, bonobos were observed avoiding visitor areas during busy times, preferring to interact when the zoo was quieter. This suggests that bonobos may feel uncomfortable with large crowds and prefer engaging with humans in less crowded settings.

Recent 2023 research of Rhesus Macaques have even shown zoo visitors as a source of enrichment^21^.

These studies together show the complexity of the visitor effect on chimpanzees and other primates, showing that it can influence a range of behaviors from aggression to social interaction. These studies also emphasize the importance of exhibit design and visitor management in promoting the welfare of captive primates.

Building on this previous research, this study aims to further explore chimpanzees in a sanctuary, with limited and guided visitor access. The following sections detail the methods used and the specific findings observed.

## 3. Results

We found a significant difference in the exhibition of affiliative behaviors (excluding grooming) of chimps in presence of visitors compared to absence of visitors. We found that the full model differed significantly from the null model (LMM: χ^2^ = 5.1432, gl=1, p < 0.05), with the presence/absence of visitors (p < 0.05) and the social group (p < 0.001) being significant variables. The absence of visitors had a positive relationship with the exhibition of affiliative behaviors (p < 0.05), and the chimpanzees from the Mutamba group performed more affiliative behaviors than the Bilinga group (p < 0.001). However, in the rest of the models, the full models didn’t differ significantly with the null models (p > 0.05), so the presence of visitors didn’t affect the exhibition of these behaviors in chimpanzees.

There were no major variations in the chimpanzees’ activity budget regarding these behaviors in the presence or absence of visitors (**Figure 2, Figure 3**). In the presence of visitors, the chimpanzees dedicated 46% of their behaviors to grooming (Mutamba: 60, Bilinga: 25), compared to 51% observed in their absence (Mutamba: 64, Bilinga: 20). Self-directed behaviors accounted for 22% of activities during visits (Mutamba: 15, Bilinga: 32), versus 20% in the absence of visitors (Mutamba: 13, Bilinga: 37). Abnormal behaviors constituted 18% of the observations in the presence of visitors (Mutamba: 6, Bilinga: 36), compared to 16% in their absence (Mutamba: 7, Bilinga: 34). Affiliative behaviors represented 13% during visits (Mutamba: 18, Bilinga: 6), similar to the 13% recorded without visitors (Mutamba: 15, Bilinga: 8). Finally, agonistic behaviors were 1% of the observations in the presence of visitors (Mutamba: 1, Bilinga: 1), compared to 0.5% in their absence (Mutamba: 0.8, Bilinga: 0.4).

**Figure 1.**
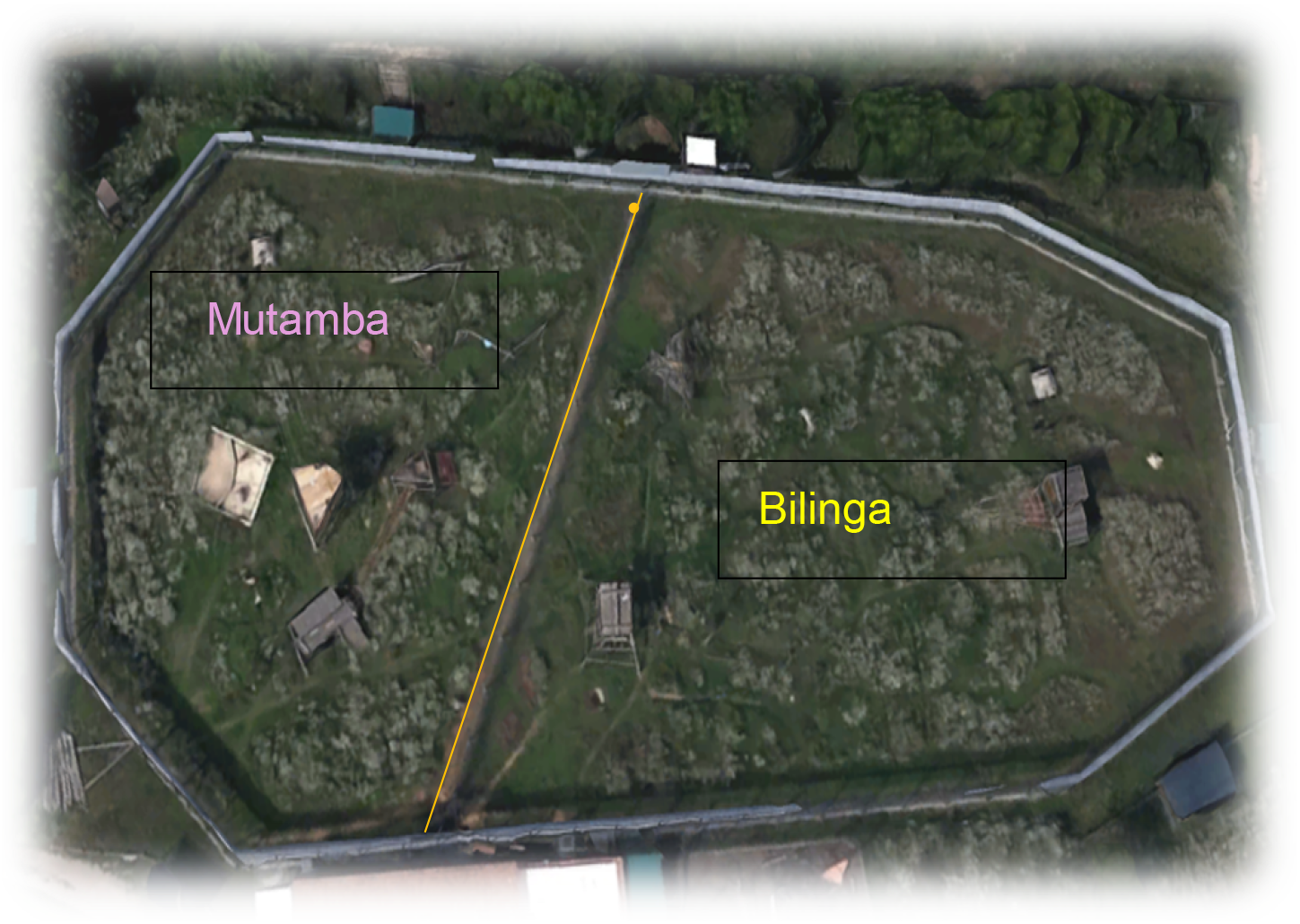
The map of outdoor enclosures shows the two groups (Mutamba and Bilinga) separated by an electric fence^29^.

**Figure 2.**
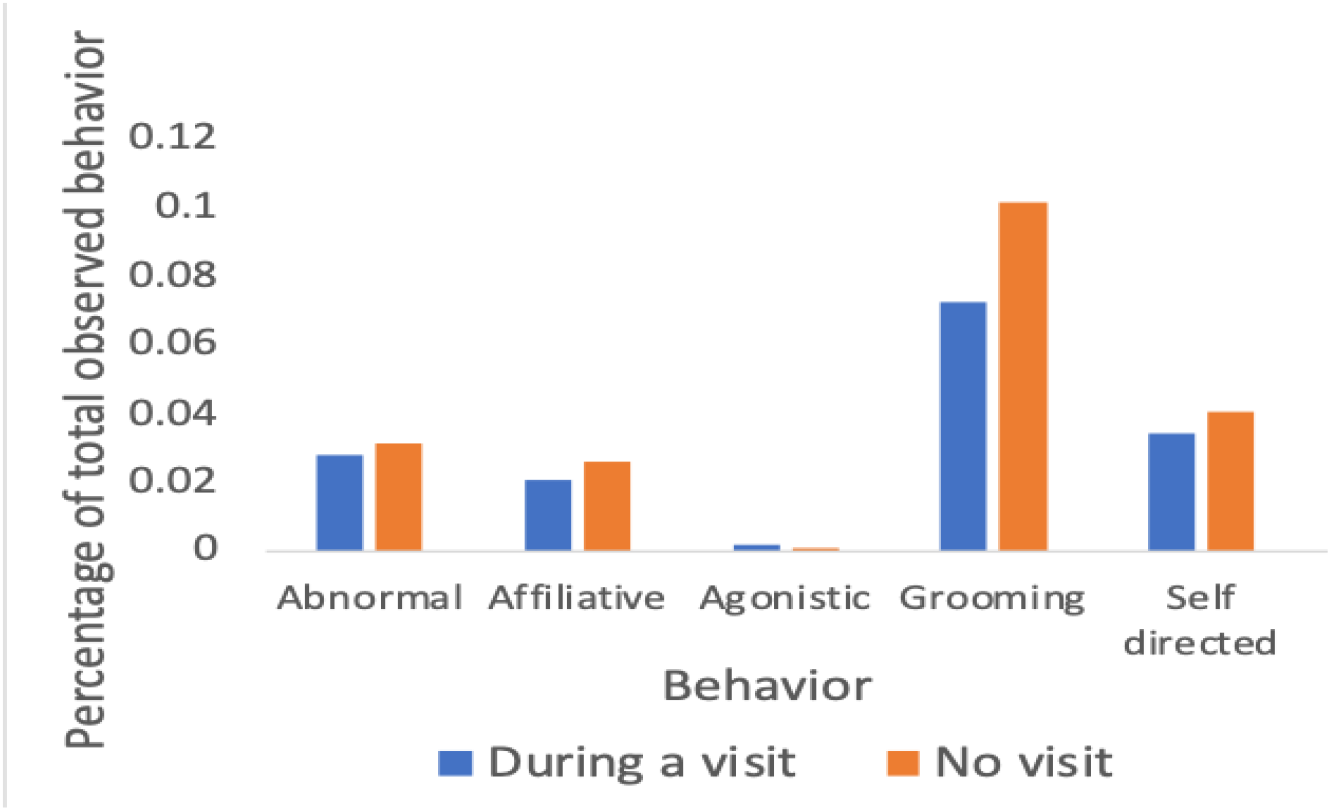
Overview of the distribution of social interactions and solitary behaviours of all individuals in the presence and absence of visitors.

**Figure 3.**
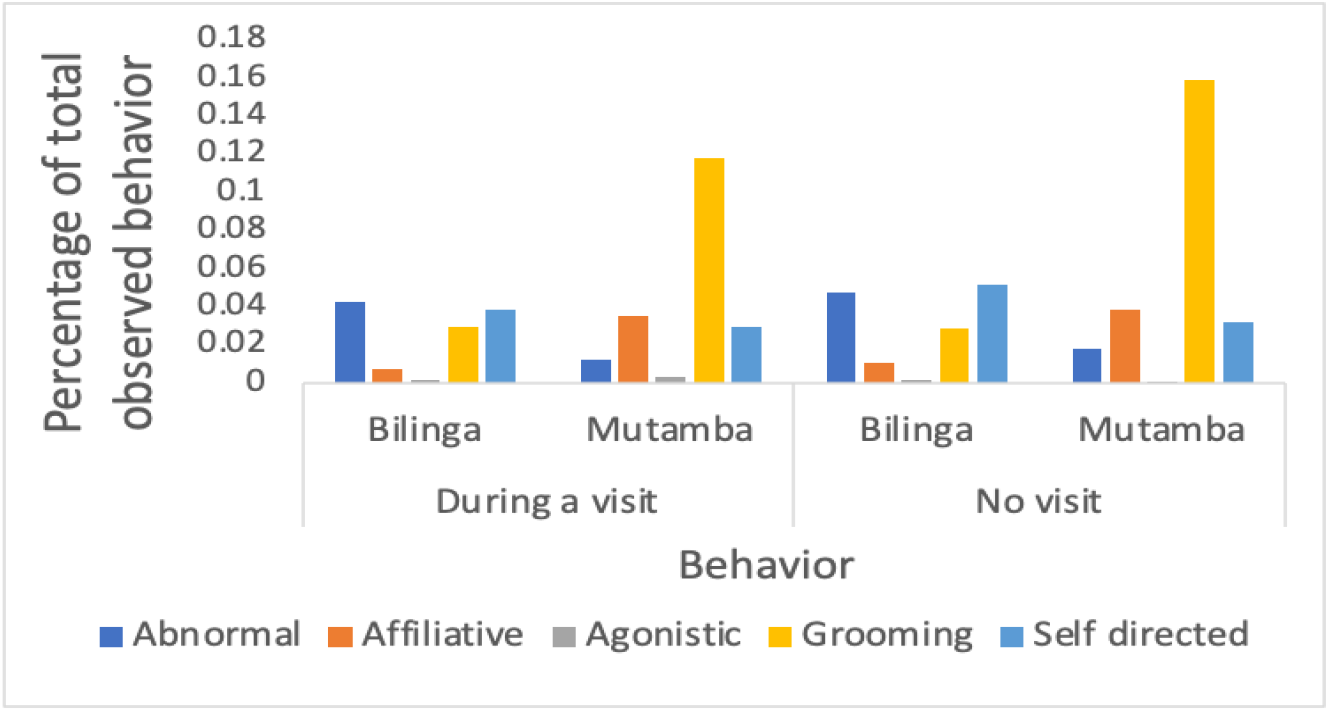
Overview of the distribution of social interactions and solitary behaviours of chimpanzees in each group in the presence and absence of visitors.

## 4. Discussion

Our results show that the visits at MONA don’t negatively affect the chimpanzees. The abnormal and self-directed behaviors are related with stress in chimps^22^. Contrary to some studies that warn of some increase in these behaviors in nonhuman primates in the presence of visitors^23,19^, we did not detect any variation in abnormal and self-directed behaviors in chimps in the presence of visitors. This may be due to the fact that visits at this rescue center are guided and limited to a maximum of 30 people per group, with the aim of preventing more than one group of visitors from being in front of the chimpanzees at the same time. Additionally, the guides prevent any interaction between the visitors and the chimpanzees. See **figure 4**.

**Figure 4.**
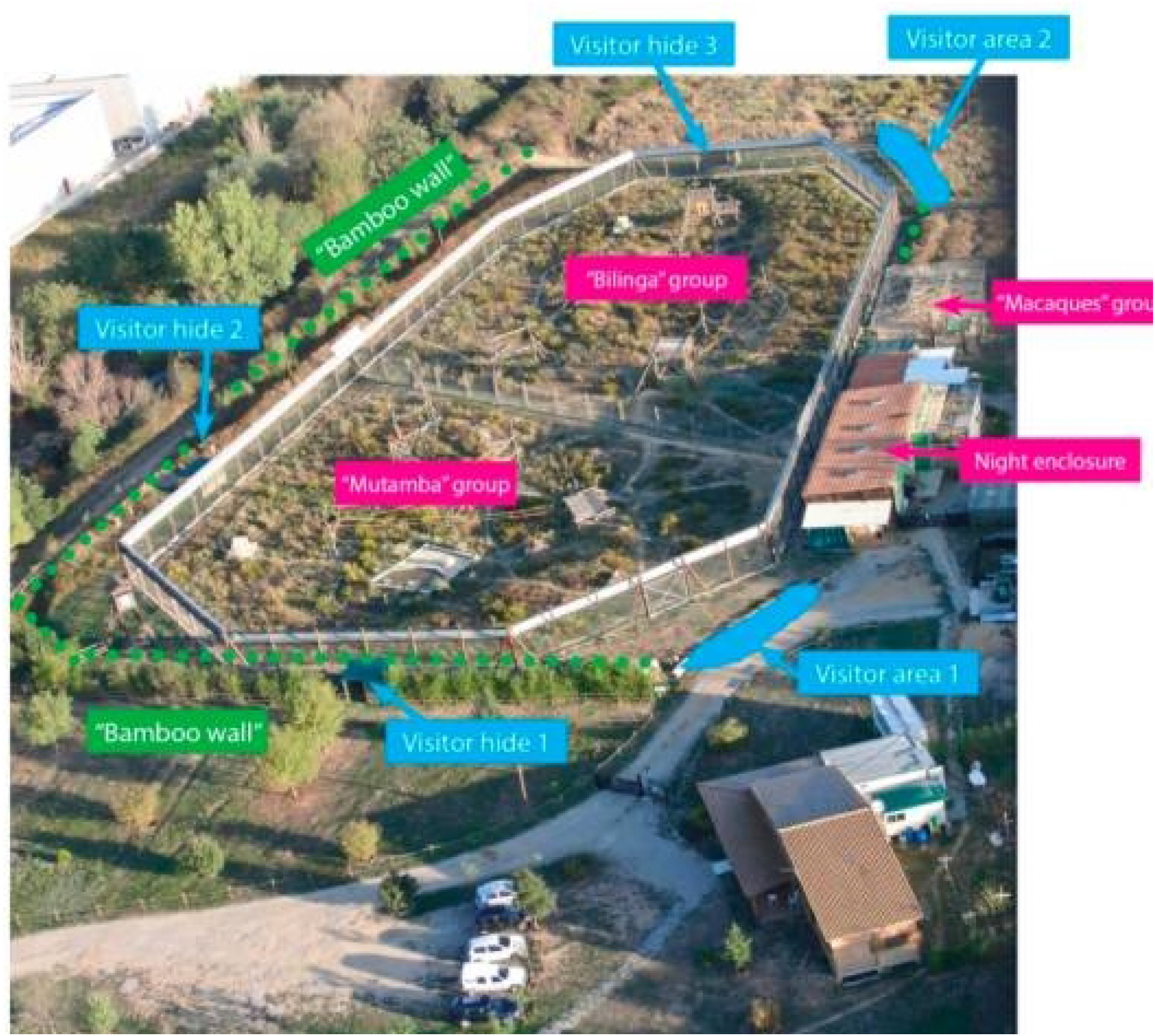
Detail of the sanctuary layout and visitor interaction system

Along with the previously mentioned factors, the visits offered at Fundació MONA aim to educate and raise awareness about the harm suffered by chimpanzees used as pets or in entertainment shows^11^. It is possible that, by becoming aware of these animals’ situations and listening to the guides’ explanations, visitors choose to adopt the most neutral behavior possible towards the chimpanzees.. Compared to other studies^24^ we did not detect differences in grooming exhibition during visits, but we did observe an increase in other affiliative behaviors recorded in our ethogram in the absence of visits. Although we have not found similar results in the literature regarding this decrease in affiliative behaviors (excluding grooming) in chimpanzees, our findings are consistent with the results of other studies on the effect of visitors on affiliative behaviors in other primate species^25^. This decrease in affiliative behaviors in the presence of visitors could be due to the chimpanzees paying more attention to the visitors than to the other members of their group. We believe that this decrease in affiliative behaviors during visits should not necessarily be seen as a negative effect. Some studies suggest that the presence of visitors can be a stimulating factor for animals in captivity^26^. Therefore, the reduced time these chimpanzees spend interacting with others in their group could be comparable to the time they would spend engaging with other enrichment provided by the caregivers at the center. This argument is supported by the lack of variation in the abnormal and self-directed behaviors exhibited by these chimpanzees in the presence and absence of visitors. On the other hand, in contrast to other studies conducted in zoos on the effect of visitors on chimpanzees, we did not detect an increase in agonistic behaviors during visits^27,28^. We believe that, due to the design of the facilities, which allows chimpanzees to hide from visitors if they wish, and given that, as we have argued previously, visitors do not represent negative stimulation but rather positive or neutral, the presence of visitors would not cause tension within the chimpanzee groups or lead to an increase in agonistic behaviors among them. Future research might focus on how visits could be structured as helpful stimuli or using AI to detect subtleties.

At Fundació MONA, we did not detect a negative effect of visits on the behavior of the chimpanzees. We observed no increase in abnormal or self-directed behaviors in the presence of visitors, nor a rise in agonistic behaviors compared to sessions without visitors. We only detected an increase in affiliative behaviors (excluding grooming) in the absence of visitors. Although this study involved a small sample of 10 chimpanzees, we believe that these guided and controlled visits are an excellent way to educate the public about chimpanzee conservation issues while also helping these centers secure funding. We conclude that limiting attendance and offering guided tours instead of open access visits to centers housing captive chimpanzees can be a good alternative for raising funds and educating the public about the species and its conservation issues, without producing a negative impact on the animals. Due to the visitor effect’s importance to animal mental health, using AI to further investigate its subtleties could be a next step.

## 5. Methods

### 5.1. Subjects and place of study

The study population consisted of 10 former pet and entertainment chimpanzees (20 to 41 years old, mean age: 29.60, SD: ±7.69) living at the primate recue and rehabilitation center Fundación MONA, located in the north of Spain, Cataluña (**Figure 1**). The chimps were separated into 2 groups: Mutamba and Bilinga, both consisting of 5 chimps, 2 females and 3 males (mean age Mutamba: 26.60, SD: ±7.36; mean age Bilinga: 32.6, SD: ±7.50). Depending on their age, animals were labeled as adults or seniors, with seniors being 35 years of age or older (only 3 males were labeled as seniors in our sample) (**Table 1**). The two groups were separated by a steel mesh and an electric fence. Their outdoor environments featured natural terrain with Mediterranean vegetation, along with various artificial climbing structures like towers, wooden platforms, bridges, ropes, and daily changing enrichment items.

**Table 1.** individual chimpanzee characteristics.

| ID | Sex | Age | (Estimated)<br>year of birth | group |
| --- | --- | --- | --- | --- |
| Africa | F | Adult | 1999 | Mutamba |
| Waty | F | Adult | 1997 | Mutamba |
| Bongo | M | Adult | 2000 | Mutamba |
| Marco | M | Senior | 1984 | Mutamba |
| Juanito | M | Adult | 2003 | Mutamba |
| Cheeta | F | Senior | 1986 | Bilinga |
| Coco | F | Adult | 1994 | Bilinga |
| Nico | M | Adult | 2001 | Bilinga |
| Tom | M | Senior | 1985 | Bilinga |
| Victor | M | Senior | 1982 | Bilinga |

### 5.2. Data collection and analysis

The data has been collected in 20-minute sessions that have been distributed uniformly throughout the day and the observation period. It was a total of 431.6 h of observation (Mutamba: 206.3 h; Bilinga: 225.3 h), spread over 6 months (January to June 2024). We coded data using instantaneous scan sampling every 2 min of all individuals in view using the ZooMonitor data scoring software^30^. During these sessions, data on social (agonistic and affiliative) and solitary (abnormal and self-directed) behaviors of the chimpanzees were collected (**Table 2**). We classified the observations into two conditions: During visits and absence of visits. We recorded 122 h of observation during visits (Mutamba: 54.7 h; Bilinga: 77.3 h), and 309.6 h of observation in the absence of visits (Mutamba: 151.6 h; Bilinga: 158 h). Data from trained observers, who passed the inter-observer reliability test (agreement ≥ 85%) with the head of research at the center (D. Crailsheim), were used for this study.

**Table 2.**
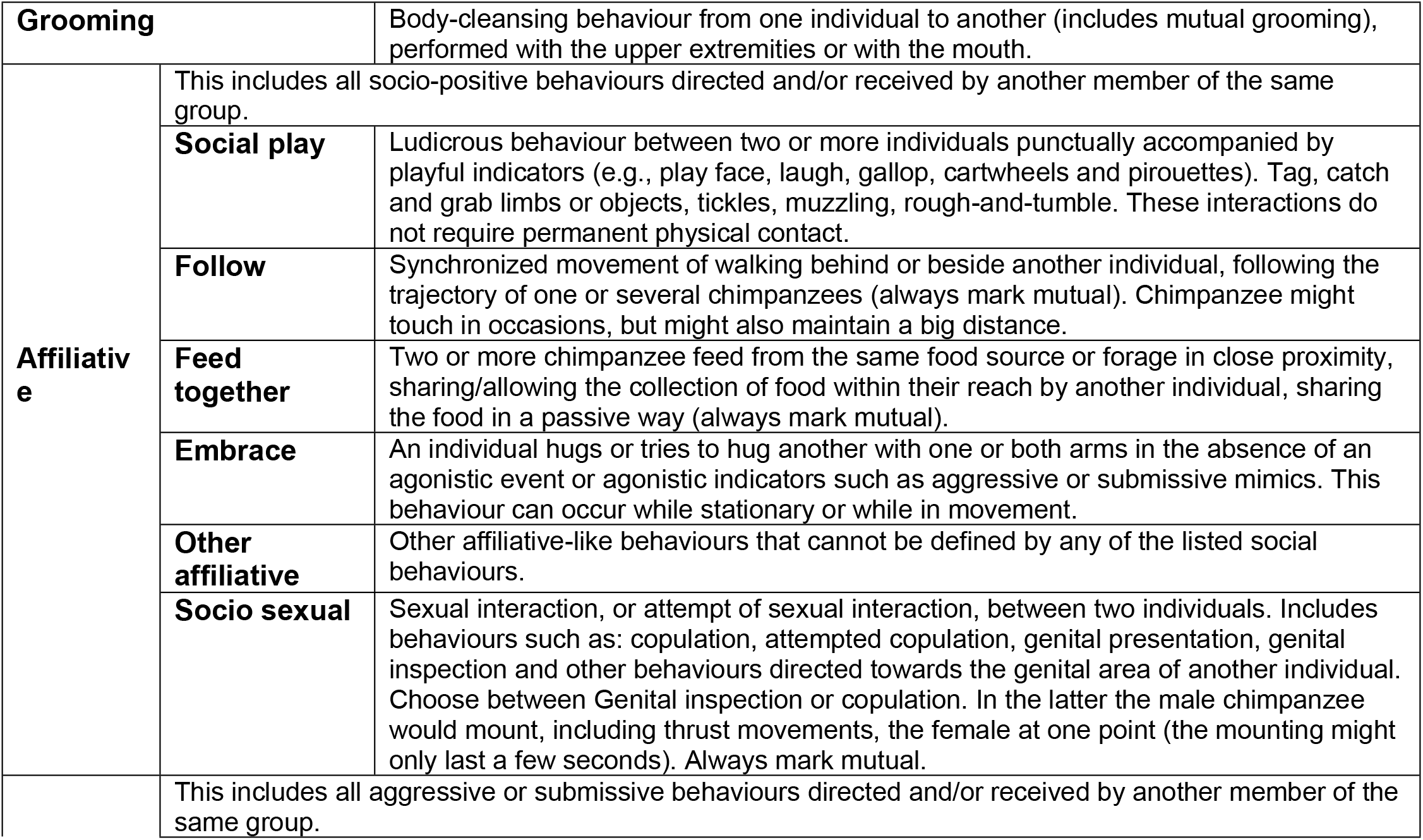
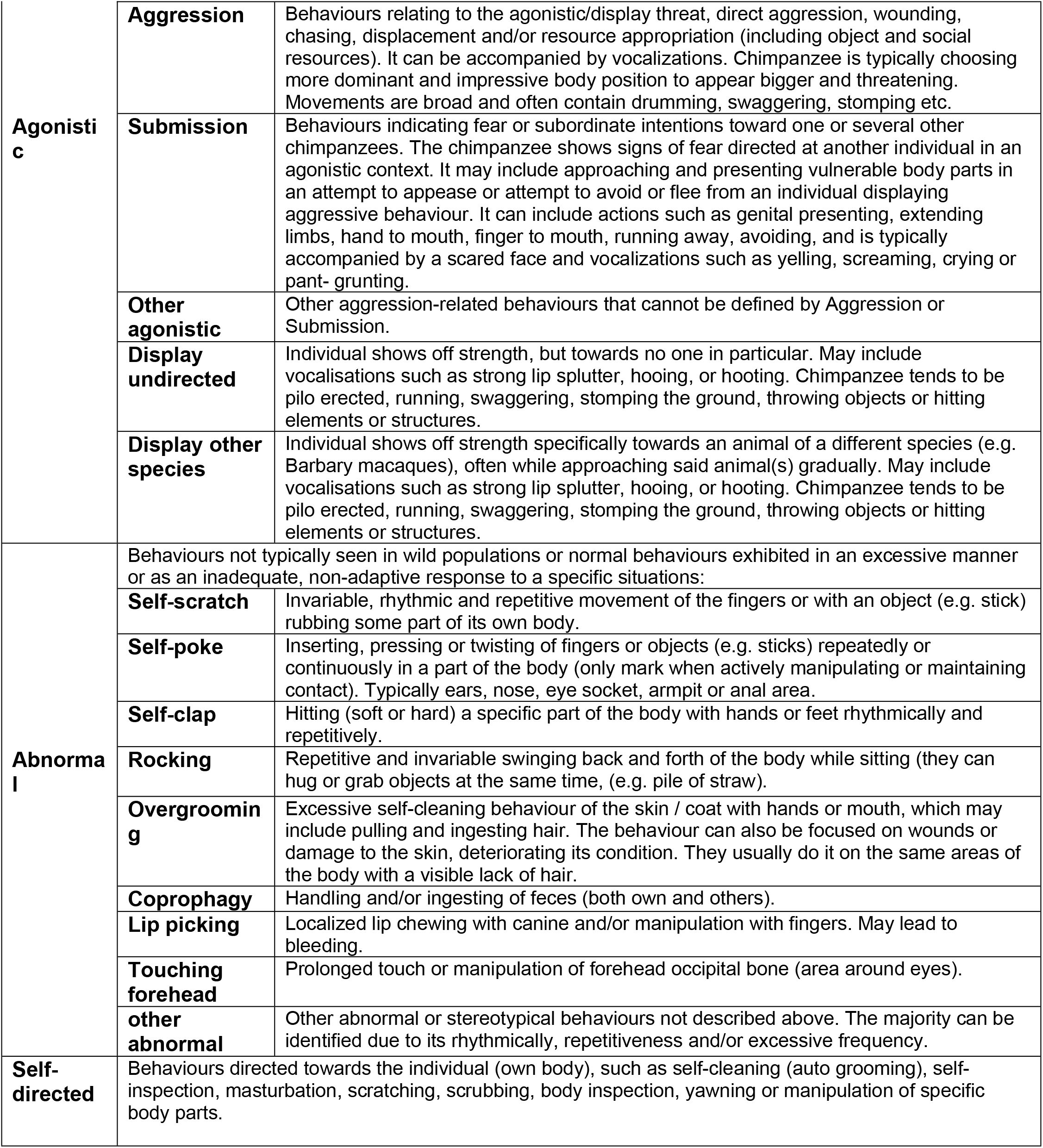
Chimpanzee ethogram behaviors.

Data was analyzed using the software program R^31^. We ran five statistical models (Grooming, Affiliative, Agonistic, Self-directed and Abnormal) to assess the impact of visits on the chimpanzees. All models had the same structural configuration, changing only the dependent variable. In each model we included sex, age, and group to which each chimpanzee belonged as control variables. Finally, we included the individual ID as a random factor.

## 6. Acknowledgements

Authors would like to thank Dr. Dietmar Crailsheim for his guidance and advice with this research. Thanks also to all the MONA researchers who helped gather the data over the period. Thanks finally to the chimpanzees for adhering well to the every two minute observation metric.

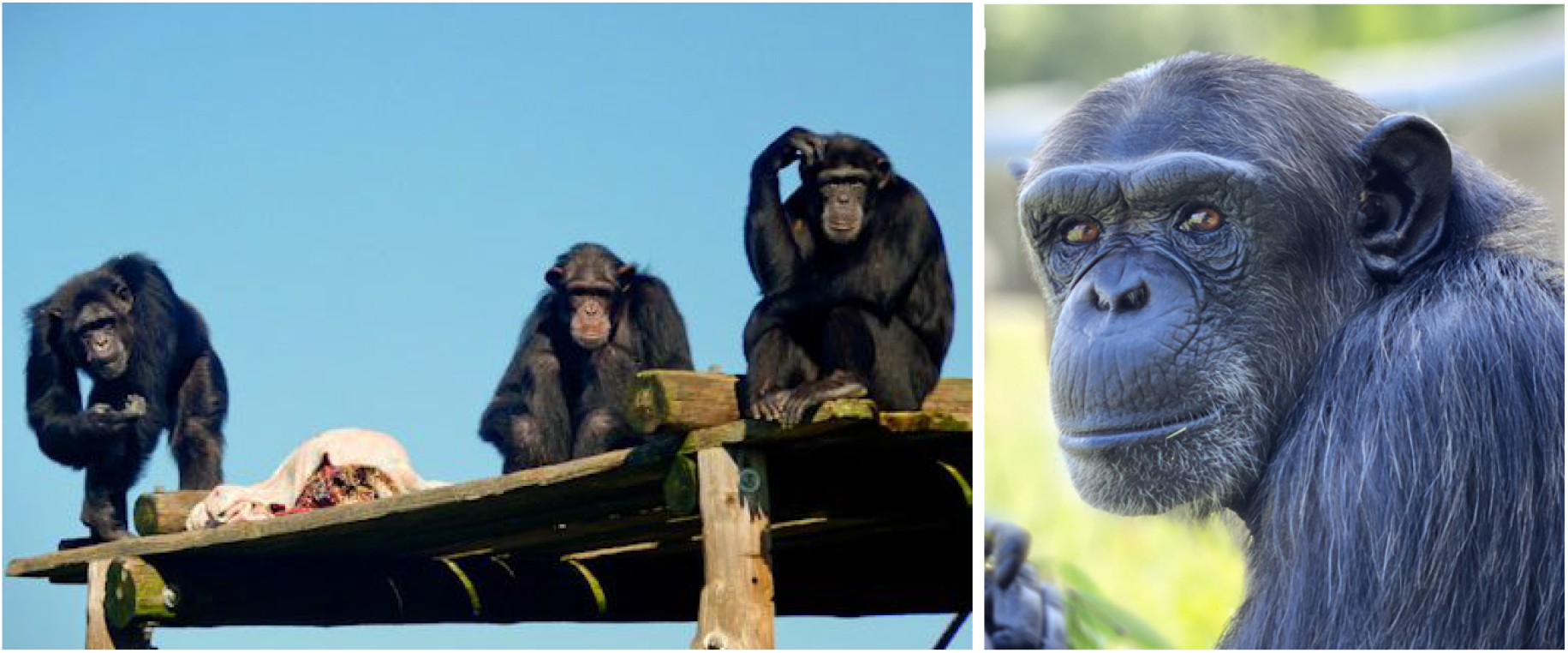

## References

(1) Aebischer, T.; Siguindo, G.; Rochat, E.; Arandjelovic, M.; Heilman, A.; Hickisch, R.; Vigilant, L.; Joost, S.; Wegmann, D. First Quantitative Survey Delineates the Distribution of Chimpanzees in the Eastern Central African Republic. Biological Conservation 2017, 213, 84–94. 10.1016/j.biocon.2017.06.031.

(2) Chivers, D. J. J. Goodall 1986. The Chimpanzees of Gombe: Patterns of Behavior. Harvard University Press, Cambridge (Massachusetts). 673 Pages. ISBN 0-674-11649-6. Price: £19.95 (Hardback). Journal of Tropical Ecology 1987, 3 (2), 190–191. 10.1017/S0266467400002029.

(3) Nishida, T. A Quarter Century of Research in the Mahale Mountains: An Overview. The Chimpanzees of the Mahale Mountains; Sexual and Life History Strategies 1990.

(4) Humle, T.; Maisels, F.; Oates, J. F.; Plumptre, A.; Williamson, E. A. Pan Troglodytes. The IUCN Red List of Threatened Species 2016: E.T15933A129038584. 10.2305/IUCN.UK.2016-2.RLTS.T15933A17964454.En. Accessed on 30 July 2024.

(5) Hicks, T. C.; Darby, L.; Hart, J.; Swinkels, J.; January, N.; Menken, S. Trade in Orphans and Bushmeat Threatens One of The Democratic Republic of the Congo’s Most Important Populations of Eastern Chimpanzees.

(6) Freeman, H. D.; Ross, S. R. The Impact of Atypical Early Histories on Pet or Performer Chimpanzees. PeerJ 2014, 2, e579. 10.7717/peerj.579.

(7) Schroepfer, K. K.; Rosati, A. G.; Chartrand, T.; Hare, B. Use of “Entertainment” Chimpanzees in Commercials Distorts Public Perception Regarding Their Conservation Status. PLOS ONE 2011, 6 (10), e26048. 10.1371/journal.pone.0026048.

(8) Anthropogenic disturbance and chimpanzee (Pan troglodytes) habitat use in the Masito-Ugalla Ecosystem, Tanzania | Journal of Mammalogy | Oxford Academic. https://academic.oup.com/jmammal/article/101/6/1660/5955477 (accessed 2024-07-31).

(9) Fultz, A. A Guide for Modern Sanctuaries with Examples from a Captive Chimpanzee Sanctuary. Animal Studies Journal 2017, 6 (2), 9–29.

(10) Llorente, M.; Riba, D.; Ballesta, S.; Feliu, O.; Rostán, C. Rehabilitation and Socialization of Chimpanzees (Pan Troglodytes) Used for Entertainment and as Pets: An 8-Year Study at Fundació Mona. Int J Primatol 2015, 36 (3), 605–624. 10.1007/s10764-015-9842-4.

(11) Feliu, O.; González-Zamora, A.; Riba, D.; Sauquet, T.; Sánchez-López, S.; Maté, C. The Impact of Sanctuary Visits on Children’s Knowledge and Attitudes toward Primate Welfare and Conservation. PeerJ 2023, 11, e15074. 10.7717/peerj.15074.

(12) van Leeuwen, E. J. C.; Bruinstroop, B. M. C.; Haun, D. B. M. Early Trauma Leaves No Social Signature in Sanctuary-Housed Chimpanzees (Pan Troglodytes). Animals 2023, 13 (1), 49. 10.3390/ani13010049.

(13) Weiss, A.; Inoue-Murayama, M.; Hong, K.-W.; Inoue, E.; Udono, T.; Ochiai, T.; Matsuzawa, T.; Hirata, S.; King, J. E. Assessing Chimpanzee Personality and Subjective Well-Being in Japan. American Journal of Primatology 2009, 71 (4), 283–292. 10.1002/ajp.20649.

(14) Bonnie, K. E.; Ang, M. Y. L.; Ross, S. R. Effects of Crowd Size on Exhibit Use by and Behavior of Chimpanzees (Pan Troglodytes) and Western Lowland Gorillas (Gorilla Gorilla) at a Zoo. Applied Animal Behaviour Science 2016, 178, 102–110. 10.1016/j.applanim.2016.03.003.

(15) Hansen, B. K.; Hopper, L. M.; Fultz, A. L.; Ross, S. R. Understanding the Behavior of Sanctuary-Housed Chimpanzees During Public Programs. Anthrozoös 2020, 33 (4), 481–495. 10.1080/08927936.2020.1771055.

(16) Tier, B. The Effect of Visitor Numbers on the Behaviour of Captive Bonobos (Pan Paniscus) at Twycross Zoo.; 2018.

(17) Bolte, J. Effects of Visitor Group Size on the Number of Abnormal Behaviors in Captive Bonobos (Pan Paniscus) Housed in Outdoor and Indoor Zoo Exhibits. Posters-at-the-Capitol 2018.

(18) Hosey, G.; Ward, S.; Melfi, V. The Effect of Visitors on the Behaviour of Zoo-Housed Primates: A Test of Four Hypotheses. Applied Animal Behaviour Science 2023, 263, 105938. 10.1016/j.applanim.2023.105938.

(19) Mallapur, A.; Sinha, A.; Waran, N. Influence of Visitor Presence on the Behaviour of Captive Lion-Tailed Macaques (Macaca Silenus) Housed in Indian Zoos. Applied Animal Behaviour Science 2005, 94 (3), 341–352. 10.1016/j.applanim.2005.02.012.

(20) Kuhar, C. W. Group Differences in Captive Gorillas’ Reaction to Large Crowds. Applied Animal Behaviour Science 2008, 110 (3), 377–385. 10.1016/j.applanim.2007.04.011.

(21) Sharma, S.; Khanal, L.; Shrestha, S.; Pandey, N.; Bellanca, R. U.; Kyes, R. C. Zoo Visitors as a Source of Enrichment to Reduce Abnormal Behavior in Captive Rhesus Macaques (<em>Macaca Mulatta</Em>) in the Central Zoo, Kathmandu, Nepal. Journal of Animal Behaviour and Biometeorology 2023, 11 (1), e2023005–e2023005. 10.31893/jabb.23005.

(22) Kutsukake, N. Assessing Relationship Quality and Social Anxiety among Wild Chimpanzees Using Self-Directed Behaviour. Behaviour 2003, 140 (8/9), 1153–1171.

(23) Animal–visitor interactions in the modern zoo: Conflicts and interventions - ScienceDirect. https://www.sciencedirect.com/science/article/abs/pii/S0168159109001890 (accessed 2024-08-31).

(24) Wood, W. Interactions among Environmental Enrichment, Viewing Crowds, and Zoo Chimpanzees (Pantroglodytes). Zoo Biology 1998, 17 (3), 211–230. 10.1002/(SICI)1098-2361(1998)17:3<211::AID-ZOO5>3.0.CO;2-C.

(25) Chamove, A. S.; Hosey, G. R.; Schaetzel, P. Visitors Excite Primates in Zoos. Zoo Biology 1988, 7 (4), 359–369. 10.1002/zoo.1430070407.

(26) Farley, A. Comparison of Chimpanzee (Pan Troglodytes) Behavior on Tour and Non-Tour Days at Chimpanzee Sanctuary Northwest. All Master’s Theses 2016.

(27) J. M. Stevens; Alonso, A. S.; Aerts, T.; Vervaecke, H. The Behaviour of a Group of Chimpanzees: Influence of Spatial Crowding and Visitor Numbers. 2008.

(28) Maki, S.; Alford, P. L.; Bramblett, C. The Effects of Unfamiliar Humans on Aggression in Captive Chimpanzee Groups. In American Journal of Primatology; Wiley-Liss Div John Wiley & Sons Inc 605 Third Ave, New York, NY 10158-0012, 1987; Vol. 12, pp 358–358.

(29) Google Earth. https://earth.google.com/web/search/fundacio+MONA/@41.90222489,2.81687456,94.17459128a,163.22913583d,35y,-78.9997068h,17.74999476t,0r/data=CigiJgokCe7JJ-qx80RAEaCjd8JJ80RAGbTGRglyjgZAIanDBIG6hwZA (accessed 2024-07-30).

(30) Wark, J. D.; Cronin, K. A.; Niemann, T.; Shender, M. A.; Horrigan, A.; Kao, A.; Ross, M. R. Monitoring the Behavior and Habitat Use of Animals to Enhance Welfare Using the ZooMonitor App. AB & C 2019, 6 (3), 158–167. 10.26451/abc.06.03.01.2019.

(31) Team, R. C. R: A Language and Environment for Statistical Computing; R Core Team: Vienna, Austria, 2022. Available at: https://www.r-project.org (accessed February 17, 2022) 2021.

